# Processing reliant on granule cells is essential for motor learning but dispensable for many cerebellar-dependent behaviors

**DOI:** 10.1101/2024.11.10.622849

**Authors:** Joon-Hyuk Lee, Chong Guo, Shuting Wu, Aliya Norton, Soobin Seo, Zhiyi Yao, Wade G. Regehr

## Abstract

Cerebellar dysfunction leads to motor, learning, emotional, and social deficits. It is assumed that these deficits arise from impaired processing of mossy fiber inputs that activate granule cells (GCs) that in turn excite Purkinje cells (PCs). However, high-frequency spontaneous PC firing might also influence behaviors. To clarify how the cerebellum regulates behaviors, we compared the effects of disrupting either GC signaling, which selectively perturbs cerebellar processing, or PC signaling, which disrupts cerebellar processing and spontaneous PC firing. We find that both GC and PC signaling are required for eyeblink conditioning and vestibulo-ocular reflex (VOR) learning. However, disrupting PC signaling impairs baseline VOR, anxiety, and social behaviors, but abolishing GC signaling does not. This establishes that cerebellar processing is essential for motor learning, but is not required for many cerebellum-dependent behaviors. This suggests that such behaviors could be rescued by elevating firing in downstream targets, as shown previously for social deficits.

## Main

The cerebellum contributes to coordination, balance, vestibular processing, motor learning, cognition, anxiety, as well as emotional and social behaviors[1-3]. The cerebellar cortex plays a significant role in cerebellar processing, with mossy fibers from various external sources exciting numerous granule cells (GCs) in the input layer. GCs in turn excite Purkinje cells (PCs), which are the sole outputs[1]. Much of what is known regarding cerebellar function is based on characterizing the influence of PCs. Abnormal PC signaling leads to deficits in basic motor function[4], motor learning[5, 6], social deficits consistent with autism spectrum disorder (ASD) symptoms[7-9], and many other behaviors[1-3]. PCs fire spontaneously at 30-150 Hz[4] in the absence of synaptic inputs[10, 11] and are also activated by climbing fiber inputs at 1-2 Hz that generate complex spikes that are followed by a brief pause in spontaneous firing [12, 13]. Although there are many contributing factors, there are two primary ways that PC signaling could influence behavior. First, PCs influence behaviors by processing MF inputs and conveying signals that flow through the cerebellar cortex. Second, spontaneous PC firing provides ongoing inhibition of neurons in the cerebellar nuclei (CbN). It is also possible that behavioral deficits could arise from inappropriate firing patterns. Thus, even though it is well established that PC-specific manipulations lead to many behavioral deficits, it is generally not known which of these mechanisms underlies how PCs influence specific behaviors.

Compared to the many established behavioral influences of the PC output layer, much less is known about how GCs in the input layer and processing within the cerebellar cortex influence most behaviors. GCs and PCs are both essential elements of cerebellar processing that are required to convey MF signals through the cerebellar cortex to influence PC outputs. It is therefore expected that eliminating either GC signaling or PC signaling would perturb behaviors that are regulated by MF activity and cerebellar processing, which is the case for gait, balance beam, and rotarod[4]. However, it is predicted that eliminating GC signaling would not impair behaviors that are regulated by mechanisms independent of GC signaling, such as spontaneous PC firing. This motivated us to compare the effects of disrupting either PC or GCs signaling to provide insight into how the cerebellar cortex influences numerous behaviors.

Disrupting PC signaling is known to prevent both conditioned eyeblink learning and vestibular ocular reflex learning[5, 6], but the importance of GCs is less clear. According to classic models, GCs are essential for motor learning[14], and cerebellar learning is a consequence of associative plasticity at GC-to-PC synapses[15-17]. However, a recent study found that impaired GC signaling delayed but did not prevent conditioned eyeblink learning, raising the possibility that eyeblink learning also occurs in the cerebellar nuclei independently of GCs[18]. The effects of disrupting GC signaling on VOR learning are unclear. Increased GC excitability does not significantly alter the changes in VOR gain during gain-down learning[19]. When GC signaling is reduced but not eliminated, VOR gain-down learning still occurs but is reduced in magnitude. However, GC signaling was not eliminated in these studies, and it has been shown that it is necessary to completely eliminate GC signaling to reveal deficits in balance beam test and gait[4]. Therefore, it is unclear whether GCs are essential for cerebellar-dependent motor learning.

Reducing PC signaling also leads to social deficits that are associated with ASD[9, 20], but it is not known if GC signaling is required for these social behaviors. Reduced GC signaling did not lead to social deficits[21], but the effects of eliminating GC signaling have not been assessed. As complete elimination of GC signaling is required to reveal the influence of GCs on gait and balance [4], the influence of GCs on social behaviors is unclear. Moreover, it is assumed that GCs are essential for social behaviors, as in a recent study that selectively deleted the ASD-related gene *SCN2A* from GCs, found that ocular reflexes were impaired similarly to those observed in ASD patients, and speculated that this disrupted GC signaling may also lead to social deficits[22, 23] (although this possibility was not tested).

Here we determine the behavioral consequences of disrupting either PC or GC signaling. We eliminate three types of calcium channels from either GCs or PCs to completely silence their synapses and find that in both cases conditioned eyeblink learning and VOR gain-down learning are eliminated. In contrast, disrupting PC signaling impairs baseline behaviors such as VOR, anxiety and social behaviors, but abolishing GC signaling does not. These studies establish that cerebellar processing and signals mediated by GCs are essential for motor learning, but are not required for many baseline behaviors. This suggests that many cerebellar-dependent behaviors could arise from mechanisms that are independent of GC signaling, such as alterations in the spontaneous firing of PCs or decreased PC-CbN signaling, and that rescuing deficits in such behaviors could simply be a matter of rescuing firing in downstream regions.

## Results

### The role of granule cell signaling in conditioned eyeblink learning

It is well established that disrupting PC signaling impairs conditioned eyeblink learning[6, 24] (**Extended Data Fig. 1**), but the effects of eliminating GC signaling are not known. We therefore tested whether disrupting GC signaling influences motor learning in the conditioned eyeblink test. We used mice in which the calcium channels that mediate synaptic transmission, CaV2.1, CaV2.2 and CaV 2.3, are conditionally eliminated using α6-cre[25] and GABAR6-cre[26] to target GCs [GC (α6) TKO mice and GC (GABRA6) TKO mice, respectively]. In our previous study, we demonstrated that this triple calcium channel knockout method can effectively block granule cell signaling without causing gross morphological changes[4]. Numerous previous studies have used these Cre lines to target GCs[4, 18, 19, 24, 27-29]. In these experiments, the unconditioned stimulus (US) was an air puff that always caused the eye to close (**Extended Data Fig. 2a**), and the conditioned stimulation (CS) was illumination with a white LED light. We measured the amplitude of eye closure by detecting the positions of the upper and lower eyelids (**Fig. 1a**, *yellow and red dots*) and the closed eyelid (**Fig. 1a**, *orange dot*) using deep learning. Control mice showed progressively larger conditioned responses (CRs) to the CS (i.e., eye closures) with training (**Fig. 1a**, *top*; **Fig. 1b-d, Supplementary Video 1**). In contrast, GC (α6) TKO mice did not show CS responses, even after many days of training (**Fig. 1a**, *bottom*; **Fig. 1b**, second row), although they showed normal US responses (**Fig. 1b**, second row, **Extended Data Fig. 2a**). The CR amplitude (peak eyelid closures during 0.4 - 0.5 s after beginning LED illumination) of control mice gradually increased to an average of 77.6% on day 8, compared to 8.1% for GC (α6) TKO mice (**Fig. 1c**, *top*). The CR probability (percentage of CR responses above 50% amplitude) of control mice increased to an average of 72.5% on day 8, compared to 8% for GC (α6) TKO mice (**Fig. 1c**, *bottom*). Similar results were seen for GC (GABRA6) TKO mice (**Fig. 1b**, bottom row, **Extended Data Fig. 2a, Extended Data Fig. 2b**). The CR amplitude on day 9 was 85.2 % for control mice compared to 1.5 % for GC (GABRA6) TKO mice (**Fig. 1d**, *top*), and the CR probability was 83.3 % and 0 % for control mice and GC (GABRA6) TKO mice, respectively (**Fig. 1d**, *bottom*). Together, these results demonstrate that disrupting GC signaling abolishes motor learning in the conditioned eyeblink test.

**Fig. 1.**
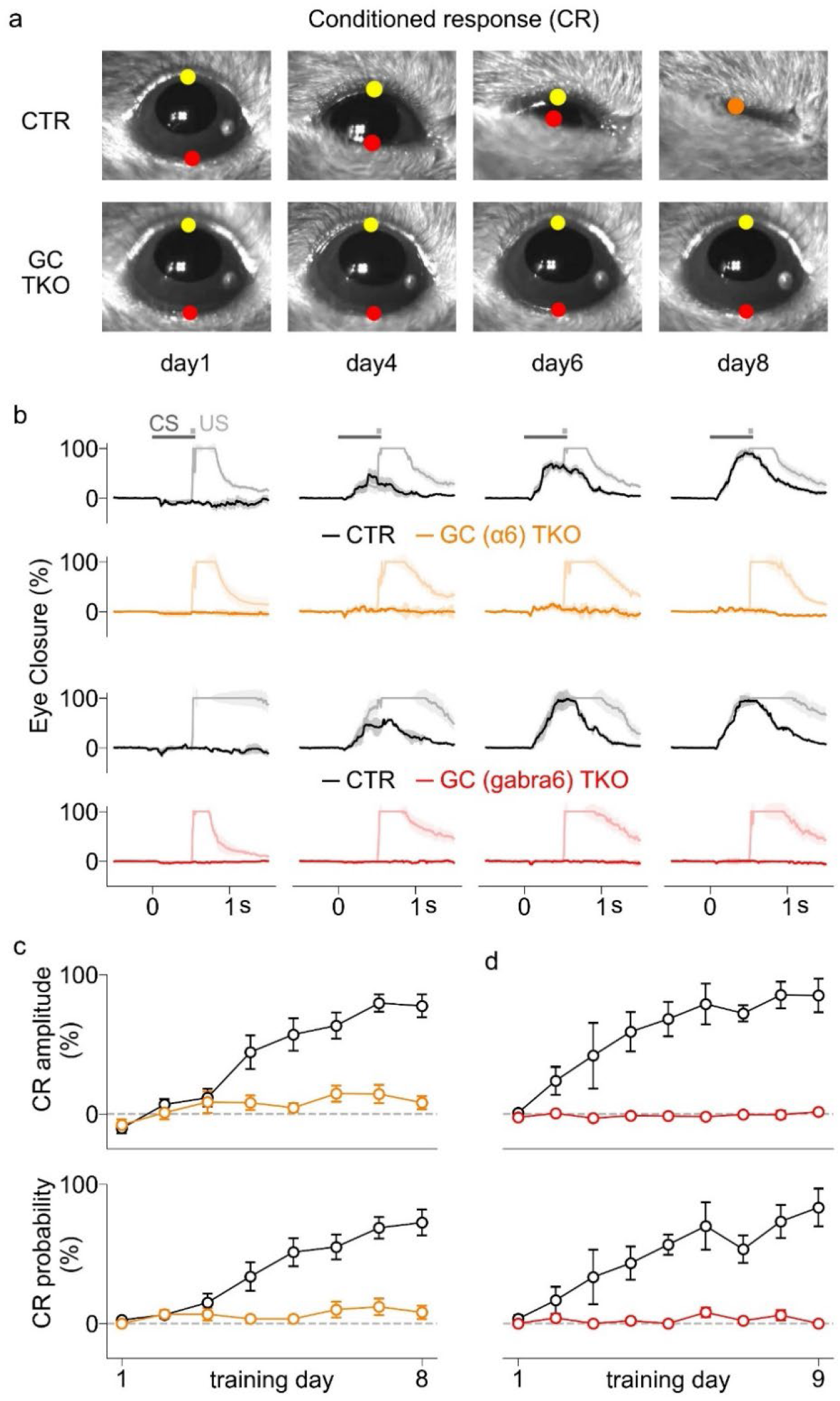
Granule cell signaling is essential for motor learning in the conditioned eye blink test. **a**, Examples of conditioned responses of a control mouse (*top row*) and a GC (α6) TKO mouse (*bottom row*) on days 1, 4, 6, and 8. The yellow and red dots indicate the positions of upper and lower eyelid, and orange dot indicates the closed eyelid, as automatically detected by deep-learning. The conditioned stimulus (CS, white LED) was presented at 0 s, and the unconditioned stimulus (US, air puff) was presented at 0.5 s. Also see Supplementary Video 1. **b**, Average eye closures are shown for GC (α6) TKO mice and corresponding control mice (first two rows), and GC (gabra6) TKO mice and corresponding control mice (third and fourth rows). CS-only trials (*dark*) and CS-US paired trials (*faint*) are plotted. The shaded area indicates SEM. **c**, Average CR amplitude (*top*) and CR probability (*bottom*) are plotted for GC (α6) TKO and control mice. **d**, As in (c) but for the GC (gaba6) TKO group and control mice. The error bars indicate SEM.

### The role of granule cell signaling in VOR learning

We also tested the extent to which GC signaling affects gain-down VOR learning. In these experiments, VOR responses were measured before and after training sessions for four consecutive days (**Fig. 2a**). During test sessions, compensatory eye movements were monitored in head-fixed mice on the turntable in the dark during vestibular stimulation (±5° turntable rotation at 0.5 Hz frequency) (**Fig. 2b**, left). During training sessions, the same vestibular stimulation was accompanied by visual stimulation (vertical bars), moving left and right in phase with vestibular stimulation ±5°, ±7.5°, ±10°, ±10° on days 1-4 respectively (**Fig. 2b**, right). There were six training sessions per day for four consecutive days. Control mice learned to reduce their compensatory eye movements and decrease their VOR gain, as shown by the progressive decrease in the angular velocity of compensatory eye movements (**Fig. 2c**, black). This learned response was apparent in all control mice tested, but was not observed in GC (α6) TKO mice (**Fig. 2d**, *orange*) and GC (GABRA6) TKO mice (**Fig. 2e**, *red*). Similarly, the PC (PCP2) TKO mice, in which PC→CbN synapses are silenced by selectively eliminating CaV2.1, CaV2.2 and CaV 2.3 from PCs, also failed to exhibit the learned response (**Fig. 2f**, *blue*)[4].

**Fig. 2.**
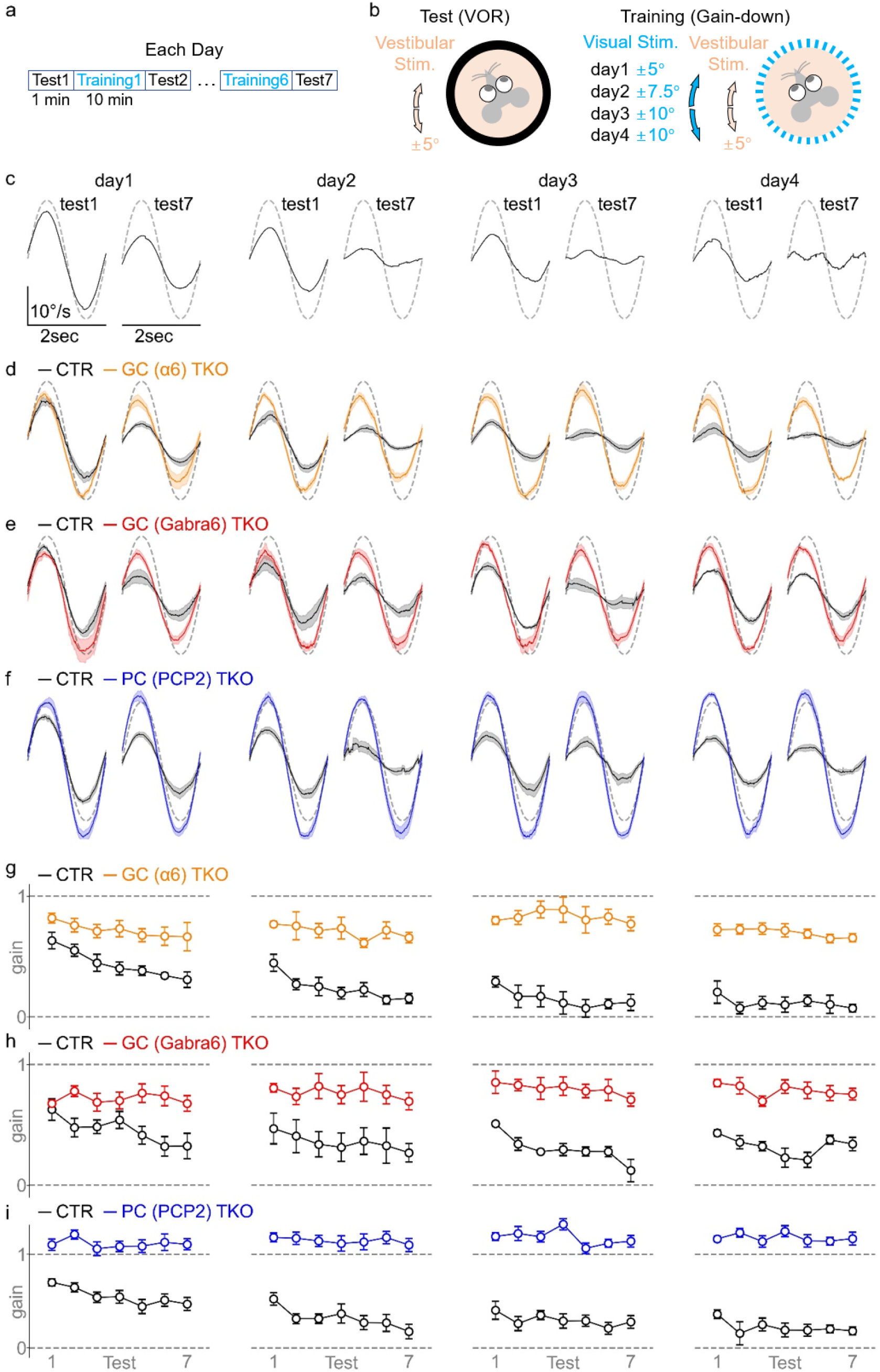
Granule cell signaling is essential for motor learning in a VOR learning test. **a**, A schematic summarizes the experimental protocol for each day in the 4-day VOR learning procedure. **b**, A schematic describes the protocols for the gain-down training (left) and the test (right). During training, vertical bars moving left and right in front of the mice are presented in phase with the vestibular stimulus (i.e. turntable rotation). During testing, a vestibular stimulus is presented in the absence of visual stimulation and the horizontal velocity of the pupil of left eye is monitored. **c**, Example angular velocities of eye movements are shown for a control mouse (black traces) along with the velocity of vestibular stimulation (dashed lines). Note that these compensatory eye movements are in opposite phase with the vestibular stimulation, but they are inverted to aid in comparison to the visual stimulation. **d**, Average eye movements are shown for GC (α6) TKO mice (colored) and control mice (black). The shaded areas indicate SEM. **e**, Same as (d) but for GC (gabra6) TKO mice. **f**, Same as (d) but for PC (PCP2) TKO mice. **g**, Changes in gain values measured in 7 tests each day for 4 days are shown for GC (α6) TKO mice (*colored*) and control mice (*black*). **h**, Same as (g), but for or GC (gabra6) TKO mice. **i**, Same as (g), but for PC (PCP2) TKO mice.

The analysis of gain values from seven tests over each of the four days revealed that GC (α6) TKO mice had impaired motor learning throughout training days (**Fig. 2g**, *orange*) compared to control mice (**Fig. 2g**, *black*). We observed similarly impaired motor learning in GC (GABRA6) TKO (**Fig. 2h**) and PC (PCP2) TKO (**Fig. 2i**), which is also evident in a graph in which all mice are plotted collectively (**Extended Data Fig. 3**). These results establish that disrupting either GC signaling or PC signaling eliminates VOR learning.

### The influence of silencing granule cell or Purkinje cell synapses on OKR, VOR and VVOR

Thus far, we have shown that disrupting GC and PC signaling leads to similar deficits in gait, rotarod, and balance beam[4], and in motor learning for both eyeblink conditioning and VOR learning (**Fig. 1, Extended Data Fig. 1, Fig. 2**). We performed additional experiments to determine if all behaviors are affected in a similar manner in GC TKO and PC TKO mice. We began by testing basic eye reflexes in the OKR test in which visual stimulation moved left and right ±5° without vestibular stimulation (**Fig. 3a**). As shown for a control mouse, we tested 0.2, 0.5, and 1 Hz stimulation (**Fig. 3b**) and compared the angular velocities of eye movements (**Fig. 3b**, left, black lines) and visual stimulation (**Fig. 3b**, left, dashed lines). These traces are replotted and normalized (**Fig. 3b**, *middle*) and the gain was determined for each frequency (**Fig. 3b**, *right*). As established previously, OKR performance is superior for low frequency visual stimulation[5, 6, 19, 27]. Gain in the OKR assay was reduced in GC (α6) TKO mice for 0.5 and 1 Hz stimulation (**Fig. 3c**), in GC (GABRA6) TKO mice for 0.5 Hz stimulation (**Fig. 3d**), and in PC (PCP2) TKO mice for 0.5 and 1 Hz stimulation (**Fig. 3e**). These findings show that disrupting GC signaling and PC signaling similarly impairs OKR performance for 0.5 and 1 Hz stimulation.

**Fig. 3.**
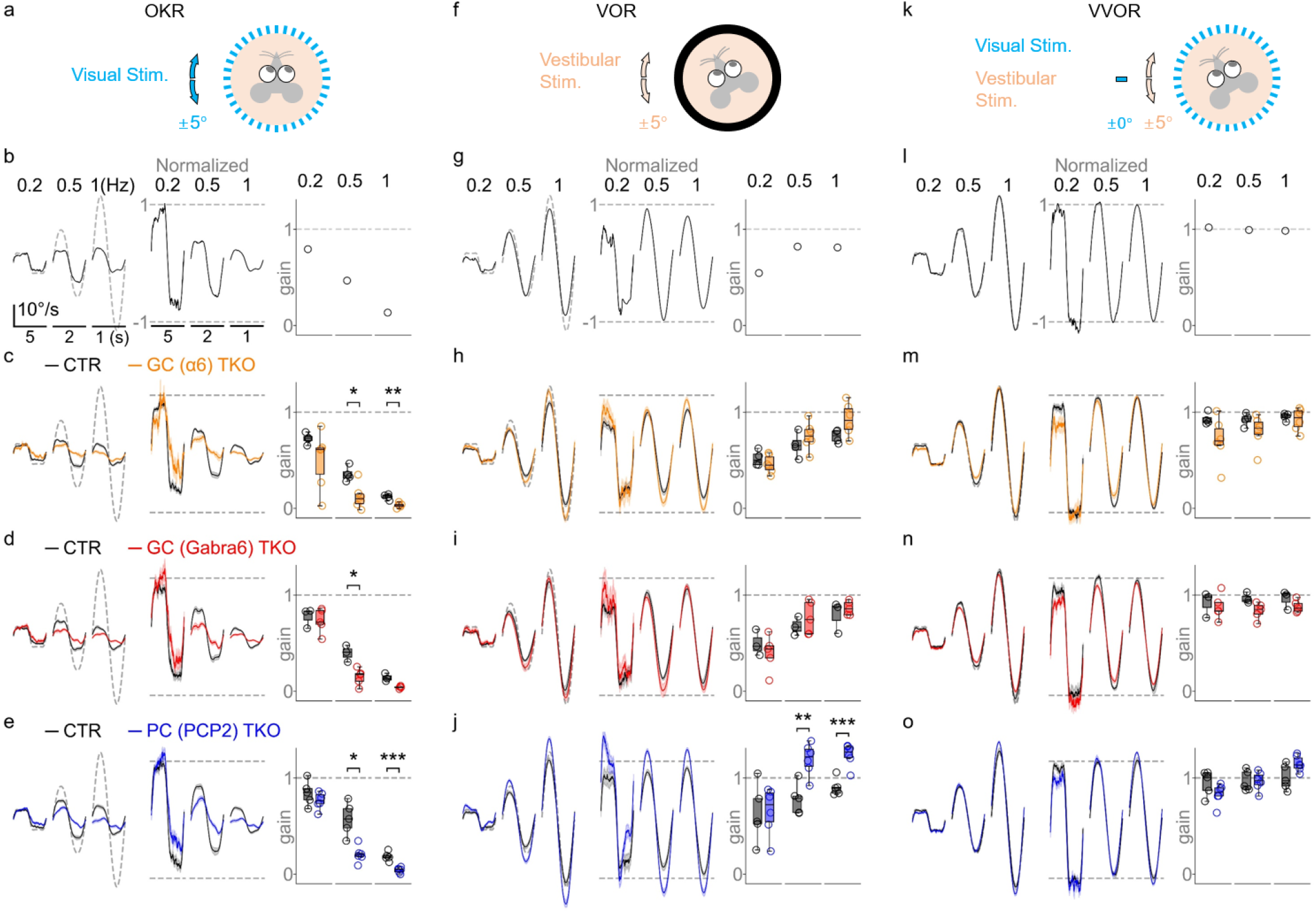
The effects of granule cell and Purkinje cell signaling on ocular reflexes. **a**, A schematic is shown for an OKR test, in which visual stimulation (vertical bars moving left and right) is presented to a head-fixed mouse. **b**, Example responses of a control mouse in the OKR test with visual stimulation at the indicated frequencies. Left, velocities of visual stimuli (gray dashed lines) and the corresponding pupil velocities (black lines) are shown. Middle, pupil velocities are replotted by normalizing to the peak velocities of visual stimulation. Right, gain values [(pupil velocity)/(peak velocity of visual stimulation)] are shown. **c**,**d**,**e**, Average OKR responses are shown for GC (a6) TKO (c), GC (gabra6) TKO (d), and PC (PCP2) TKO (e) mice. The solid lines and shaded areas (left and middle) indicate mean and SEM. Each circle (right) indicates a mouse. **f**, A schematic is shown for a VOR test, in which vestibular stimulation (turntable rotation) is presented in darkness. **g**, Same as in (b) but for the VOR experiment with the dashed lines corresponding to vestibular stimulation. Pupil velocities in (g-j) are inverted for display purposes. **h**,**i**,**j**, Same as in (c,d,e) but for the VOR test. **k**, A schematic is shown for a VVOR test, in which vestibular stimulus is presented as in (f), but in combination with stationary visual stimulation. **l**, Same as in (g) but for the VVOR test. Pupil velocities in (l-o) are inverted for display purposes. **m**,**n**,**o**, Same as in (c,d,e) but for the VVOR test.

In the VOR test, vestibular stimulation was ±5° turntable rotation in the dark (**Fig. 3f**). As shown for a control mouse, we tested 0.2, 0.5, and 1 Hz vestibular stimulation (**Fig. 3g**) and compared the angular velocities of eye movements (**Fig. 3g**, left, black lines) with the angular velocity of vestibular stimulation (**Fig. 3g**, left, dashed lines). These traces are normalized and replotted (**Fig. 3g**, *middle*), and the gain was determined for each frequency (**Fig. 3g**, *right*). The gain of compensatory eye movements is nearly 1 for 0.5 and 1 Hz stimulation, and lower for 0.2 Hz stimulation. The properties of VOR were unchanged in GC (α6) TKO mice (**Fig. 3h)** and GC (GABRA6) TKO mice (**Fig. 3i)**. PC (PCP2) TKO mice exhibited an abnormal VOR response at 0.5 and 1 Hz stimulation in which the angular velocities of eye movements were larger than expected (**Fig. 3j**). Close inspection of the 0.2Hz VOR responses also revealed a transient abnormal VOR response for 0.2 Hz stimulation in PC (PCP2) TKO mice (mean ± sem, CTR, 0.74 ± 0.12, TKO, 1.22 ± 0.11; t-test, t(8.7)=2.9, P=0.017), but not in GC (α6) TKO mice (CTR, 0.67 ± 0.05, TKO, 0.82 ± 0.08; t-test, t(7.5)=1.5, P=0.183) or GC (GABRA6) TKO mice (CTR, 0.65 ± 0.03, TKO, 0.67 ± 0.15; Mann-Whitney test, U=6.0, P=0.786). These findings establish that disrupting GC signaling and PC signaling does not always influence behaviors in the same way.

The VVOR test is similar to the VOR test, with the exception that experiments are performed with stationary visual stimulation (**Fig. 3k**). The addition of visual stimulation greatly improves the performance of compensatory eye movements, and, as shown for a control mouse, gain values at all frequencies were approximately one (**Fig. 3l**). VVOR performance was not impaired in GC (α6) TKO mice (**Fig. 3m)**, GC (GABRA6) TKO mice (**Fig. 3n)** or PC (PCP2) TKO mice (**Fig. 3o**). These findings indicate that VVOR does not require the cerebellar cortex.

### Influences of GCs and PCs on spontaneous behavior, locomotion and anxiety

We then investigated how disrupting GC signaling and PC signaling affects spontaneous behavior by using the behavioral analysis platform Motion Sequencing (MoSeq) in which the movements of individual mice within a circular arena are monitored with a depth camera, and categorized as a series of distinct three-dimensional behavioral syllables that make up spontaneous behaviors (**Fig. 4a**)[30]. MoSeq has proven to be a useful tool to detect subtle variations in behavior in mouse models[31, 32]. Usages of some syllables were strongly reduced in all TKO mice compared to control mice (**Fig. 4b**). These syllables were all related to the initiation and cessation of locomotion (**Supplementary Video 3**). The similarity in the disruption of these behaviors for GC (α6) TKO, GC (GABRA6) TKO and PC (PCP2) TKO was not surprising in light of the observation that these mice show similar deficits in gait, balance, rotarod, and OKR (**Fig. 3c-d**)[4]. We also found that the usage of some syllables was unaltered in GC (α6) TKO mice and GC (GABRA6) TKO mice, but was selectively altered in PC (PCP2) TKO mice (**Fig. 4c**). These syllables are associated with swaying behavior in a stationary state and with the initiation of motion (**Supplementary Video 3**). Although the functional importance of these behavioral differences is unclear, this unbiased analysis of spontaneous behaviors revealed the important result that disrupting PC signaling and GC signaling differentially alters some spontaneous behaviors.

**Fig. 4.**
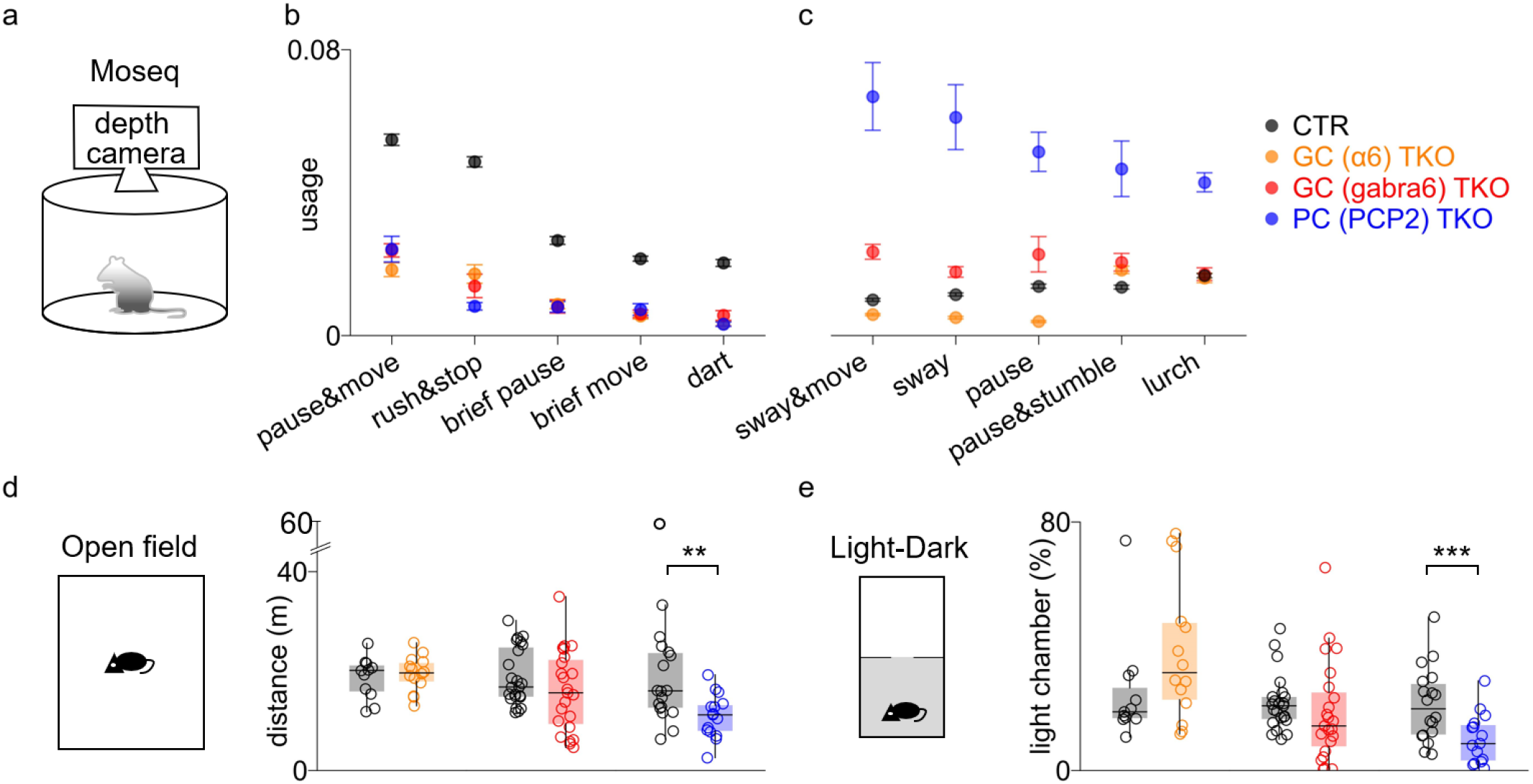
Disrupting granule cell and Purkinje cell signaling differentially influences some behaviors. **a**, A schematic shows the Moseq assay in which the 3D motion of mice is recorded by a depth camera. **b**, Moseq behavioral syllables that are observed less frequently in all TKO mice. **c**, Syllables that are most frequently observed in PC (PCP2) TKO mice (c). **d**, A schematic of open field test. **e**, The distances travelled for control (*black*), GC (a6) TKO (*orange*), GC (gabra6) TKO (*red*) and PC (PCP2) TKO (*blue*) mice are shown. Each symbol indicates a mouse. **f**, A schematic of light-dark test. **g**, The time spent in the light chamber is shown for each group.

Subsequent experiments revealed that disrupting PC and GC signaling differentially influenced additional behaviors. In the open field test, locomotion was only reduced in PC (PCP2) TKO mice and was normal in GC (α6) TKO, GC (GABRA6) TKO mice (**Fig. 4d**). Further analysis of the speed in open file test revealed that PC (PCP2) TKO mice spent more time in lower speeds (0-5 cm/s) and less time in high speed (20-100 cm/s), whereas GC TKO mice did not show differences compared to control mice (**Extended Data Fig. 4b-d**). This was surprising because PC TKO mice and GC TKO mice show similar impairments in gait and balance beam. The time spent in the center zone was unaltered in any TKO mouse line (**Extended Data Fig. 4a**). We also used a light/dark test to assess anxiety. Mice are naturally averse to bright light and tend to avoid a brightly illuminated chamber in favor of a dark chamber. The decreased time spent in the bright chamber is generally interpreted as reflecting an increase in anxiety-like behavior. We found that PC (PCP2) TKO mice spent less time in the illuminated chamber than littermate controls, but GC (α6) TKO and GC (GABRA6) TKO mice spend a comparable amount of time as littermate controls (**Fig. 4e**). These findings suggest that disrupting PC signaling increases anxiety, but disrupting GC signaling does not influence anxiety. Together, these findings reinforce the conclusion that disrupting PC signaling and GC signaling differentially affects many behaviors, including VOR, spontaneous behaviors, locomotion and anxiety.

### Granule cell signaling is not required for social behavior

We investigated how disrupting GC and PC signaling influences social behavior by using the well-established three-chamber assay in which the time spent near a social stimulus and an object are compared (**Fig. 5b**). Mice normally prefer to investigate a novel mouse rather than a novel object, as is the case for control mice in our experiments (**Fig. 5a, c**). Remarkably, in stark contrast to their severe deficits in motor functions [4] (**Fig. 1 and 2**), GC (α6) TKO mice exhibited normal social preference, and spent more time near the social stimulus than the object stimulus, similar to control mice (**Fig. 5a, c**). Social preference was also intact in GC (GABRA6) TKO mice, although they also spent more time in the middle chamber (**Extended Data Fig. 5a, 6b**). Social preferences were intact for both male and female GC (α6) TKO and GC (GABRA6) TKO mice (**Extended Data Fig. 6b**). As anticipated, PC (PCP2) TKO mice did not show a preference to social stimulus (**Fig. 5a, c**). This illustrates the ability of the synaptic silencing approach we use to reveal social deficits. Whereas more minor disruptions of PC signaling just seem to eliminate preferences for novel mice[7-9], we found that complete suppression of PC synapses also caused mice to spend extended periods of time immobile near the edge of the enclosure.

**Fig. 5.**
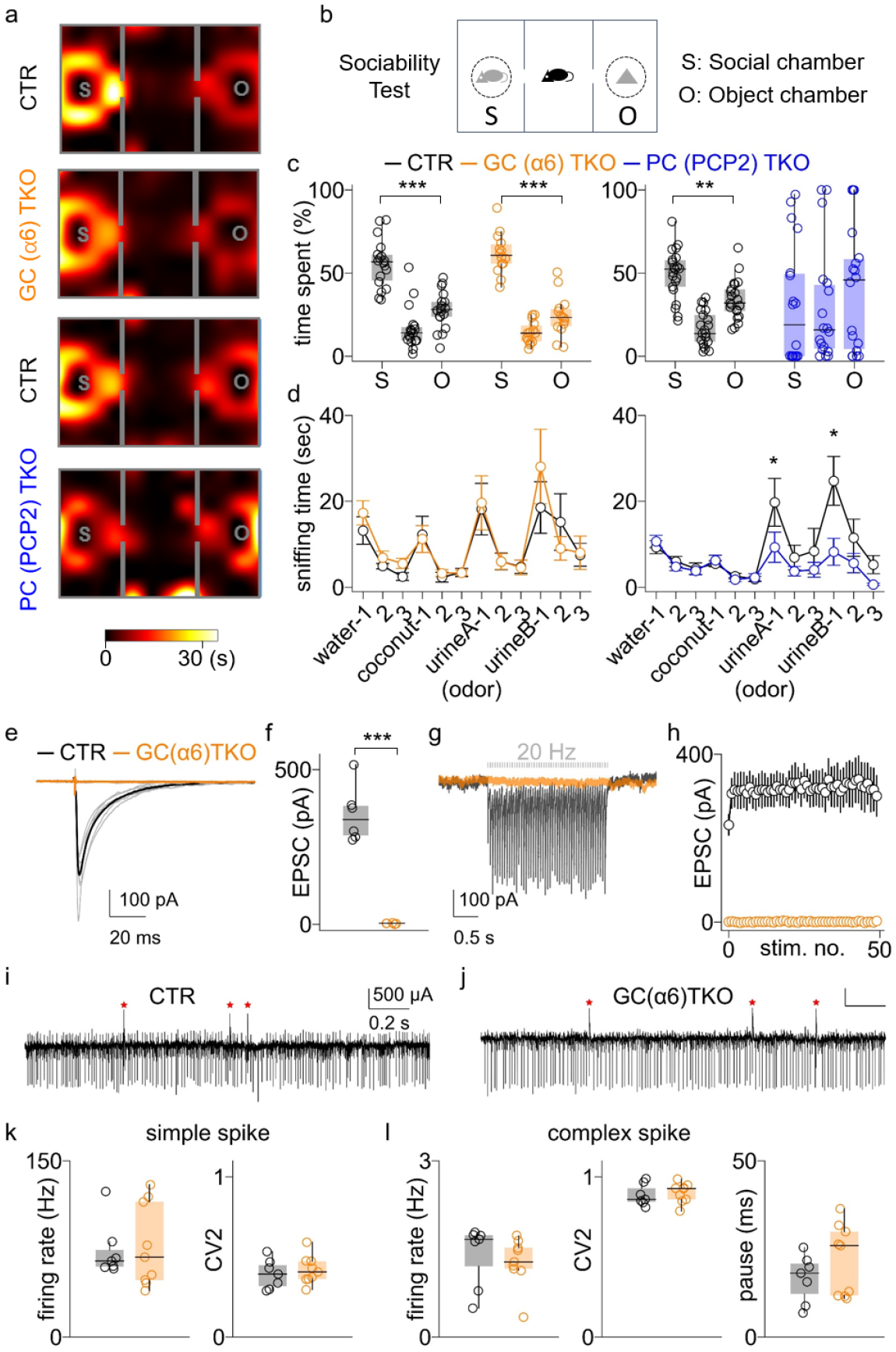
Granule cell signaling is dispensable for social behavior. **a**, Heat maps show the time spent in the chambers during the sociability test. **b**, A schematic of the sociability test. **c**, The time spent in social chamber (*S*) containing a stimulus mouse, middle chamber and object chamber (*O*) are shown for control (*black*), GC (a6) TKO (*orange*) and PC (PCP2) TKO (*blue*) mice. Mice did not show preferences for the social chamber before we introduced the stimuli (**Extended Data Fig. 6a**). Each circle indicates a mouse. **d**, The sniffing time observed in the olfactory test is summarized. Neutral cues (water, coconut) and social cues (urine of same sex - urineA, opposite sex - urineB) were presented three times consecutively. **e-h**, GC-PC EPSCs were studied in brain slices from Crus1 using whole-cell recordings. **e**, GC-PC EPSCs of GC (a6) TKO (*orange*) and control (*black*) mice are shown. Responses from each cell (thin traces) and average responses (bold traces) are shown. **f**, Peak responses are summarized. Each circle is a cell. **g**, Same as in (e), but for 20 Hz train stimulation. **h**, Peak responses to 20 Hz stimulation are summarized. Error bars indicate SEM. **i-l**, PC firing in Crus1 *in vivo* was examined in adult mice for GC (a6) TKO and control mice. **i**,**j**, The *in-vivo* PC firings in Crus1 were examined in adult mice. **i**, Simple spikes (downward spikes) and complex spikes (upward spikes marked as red asterisks) are apparent in a PC recording from a control mouse. **j**, As in i, but for GC (a6) TKO (j) mouse. **k**, Firing rates and variability (CV2) of control (*black*) and GC (a6) TKO (*orange*) mice are summarized for simple spikes. Each circle indicates a cell. **l**, As in k but for complex spikes. The durations of pauses in simple spike firing following complex spikes are also summarized.

To further assess social preference in GC (α6) TKO mice, we performed an olfactory test, measuring sniffing time in response to both neutral and social odors. Genetic manipulations that target PCs have been shown to decrease the time spent near a novel animal and impair the detection of social odors[9, 33, 34]. GC (α6) TKO mice showed normal sniffing responses to both neutral and social odors (**Fig. 5d**, left). PC (PCP2) TKO mice showed normal responses to neutral odors but significantly reduced responses to social odors (**Fig. 5d**, right), as is the case for other mouse models in which PCs are selectively manipulated and social preference is impaired[7-9, 34].

One possible explanation for the normal social behavior observed in GC (α6) TKO mice was incomplete disruption of GC signaling in Crus I, which is the region of the cerebellar cortex involved in social behaviors [7, 8]. We previously showed that synaptic transmission was eliminated at GC-PC synapses in the vermis of GC (GABRA6) TKO mice[4], but diverse types of PCs and GCs are present in different regions[35]. We therefore tested the possibility that the normal social behavior observed in GC (α6) TKO mice was due to incomplete disruption of GC signaling in Crus I. We used whole-cell electrophysiology to record GC-PC EPSCs in acute cerebellar slices containing Crus I from 8-week-old adult mice. We reliably evoked GC-EPSCs in Crus I of control mice, but no responses were observed in GC (α6) TKO mice (**Fig. 5e, f**). We also examined the possibility that prolonged high-frequency stimulation might reveal residual synaptic transmission. We stimulated GC inputs with trains of 50 stimuli and evoked sustained responses in control mice, but again did not observe any response in GC (α6) TKO mice (**Fig. 5g, h**). Synaptic transmission was also eliminated in Crus I of GC (GABRA6) TKO mice (**Extended Data Fig. 5c-e**). These results establish that intact social behaviors in GC TKO mice do not reflect incomplete disruption of GC signaling.

Having shown that signal flow through the cerebellar granule cell layer is eliminated in GC TKO mice, we examined whether disrupting GC signaling alters PC spontaneous firing in Crus I of GC (α6) TKO mice. We used multi-electrodes to record firing in Crus I PCs in awake mice. PCs fire two different types of spikes that are readily detected in extracellular recordings: simple spikes that are typical sodium-based action potentials that are present at high rates, and complex spikes that are present at 1-2 spikes/s and are evoked by spontaneous firing of the cells in the inferior olive that give rise to climbing fiber synapses (**Fig. 5i**). Simple spikes and complex spikes were also observed in GC (α6) TKO mice (**Fig. 5j**). The firing rate and variability of simple spikes (**Fig. 5k**) and complex spikes (**Fig. 5l**, left and middle) were not altered in GC (α6) TKO mice compared to littermate control mice. Complex spikes are followed by a pause in simple spike firing, and we found that the durations of such pauses following complex spikes were also unchanged (**Fig. 5l**, right). These findings indicate that disrupting GC signaling does not alter the spontaneous PC firing, and there are no compensatory changes in either PC firing or in the firing of inferior olivary neurons that give rise to climbing fibers.

## Discussion

A comparison of the behavioral effects of eliminating either GC or PC signaling revealed that the cerebellum regulates some behaviors through an unexpected mechanism. GC and PC signaling are both essential for motor learning and OKR, as shown previously for gait, balance beam, and rotarod performance[4]. This establishes that GCs and cerebellar processing are required for these behaviors. In contrast, PC signaling is essential for normal VOR, many spontaneous behaviors, normal anxiety levels, and social interactions, but GC signaling is not required for these behaviors. This indicates that cerebellar processing that relies on GC signaling is not required for these behaviors, and suggests that these behaviors are regulated by ongoing PC firing.

### The behavioral influence of PCs

Completely silencing PC synapses allowed us to attribute behavioral deficits in PC TKO mice to the lack of PC signaling[4], and the magnitude of the effects is expected to be larger than with partial suppression of PC signaling. This approach differs from most previous studies that examined the behavioral roles of PCs that were not designed to completely eliminate PC signaling. Numerous studies have determined the behavioral effects arising from selective PC targeting of genes implicated in movement disorders and autism spectrum disorder (for review, see Table 3 of [36]). These studies established the importance of PC signaling for movement and social disorders, but the observed behavioral effects could arise from decreased PC signaling or inappropriate PC firing.

PCs are known to be crucial to conditioned eyeblink learning[6, 24], but there have been conflicting reports on whether PCs play an essential role in VOR learning. Some studies have shown that introducing inappropriate firing patterns in PCs impaired gain changes in gain-down learning, but did not eliminate the changes entirely[5, 19], whereas another study reported that VOR learning was minimal[6]. Our results show that completely eliminating PC signaling abolished VOR gain-down learning (**Fig. 2f, i**), supporting the idea that PCs are essential for VOR learning.

PC-TKO mice showed a large decrease in OKR gain (reduced to 35 % of control; at 0.5 Hz CTR: 0.57, PC TKO: 0.20) (**Fig. 3e**), an increase in VOR gain with a prominent overshoot (increased to 154 % of control; at 0.5 Hz CTR: 0.79, PC TKO: 1.22, **Fig. 3j**) and unaltered VVOR gain (**Fig. 3o**). Our findings (**Fig. 3e, o**) are consistent with a previous study that suppressed PC firing by acute injection of the GABA_A_-receptor agonist, muscimol, into the flocculus that reduced OKR gain from 0.84 to less than 0.5 (0.5 Hz stimulation) and minimally affected VVOR [37]. Thus our genetic suppression of PC signaling has similar effects on OKR and VVOR as those arising from acute silencing of PCs in the monkey flocculus.

Social approach and social olfaction were strongly impaired in PC TKO mice, as in many previous studies in which PCs were manipulated[9, 33, 34, 38, 39], or PCs in subregions perturbed[7, 8]. Surprisingly, PC TKO mice also spent prolonged periods immobile next to a wall. Importantly, prolonged periods of immobility are not a secondary consequence of impaired gait, because GC (α6) mice are also ataxic but do not exhibit this behavior. Anxiety is also elevated by optogenetic activation of molecular layer interneurons in lobule VII of the vermis[3], whereas inhibition of PCs in right Crus I led to social deficits[7, 8]. Thus, it is likely that in PC TKO mice social deficits and increases in anxiety arise from disrupting PC signaling in right Crus I and lobule VII, respectively.

### GCs are essential for motor learning but are not required for many behaviors

We found that eliminating GC signaling prevents motor learning in the conditioned eyeblink test (**Fig. 1**) and prevents changes in gain down VOR learning (**Fig. 2**), which is consistent with classic models of cerebellar plasticity in which cerebellar learning arises from modifications of GC-PC synapses. Complete silencing of GC signaling was an essential element of our studies that allowed us to assess the full influence of GCs on behavior. We have previously shown that it is necessary to eliminate all calcium channels that mediate transmission at GC synapse (CaV2.1, CaV2.2 and CaV2.3) to impair gait and to strongly impair balance[4], as is the case for the GC (α6) TKO and GC (GABRA6) TKO mice we use here. Consequently, manipulations that suppress but do not completely eliminate GC signaling may provide important insights into the behavioral importance of GC signaling, but they do not reveal the full influence of GCs on behavior. This could account for the modest influence of GC-specific manipulations on behavior observed in studies that suppressed but not eliminate GC signaling. In a previous study in which inhibitory DREADDs were expressed in GCs, conditioned eyeblink learning was delayed but not eliminated, and it was concluded that slow onset learning was mediated by a mechanism within the CbN that did not involve GCs[18]. However, DREADDs can leave 30-50% of residual activity intact *in vivo* [40-42] and it is highly unlikely that inhibitory DREADDs eliminate GC firing. As we do not see any learning in our experiments when we fully eliminate GC signaling (**Fig. 2d, e, g, h**), and see no indication of a mechanism that is independent of GC signaling, it is possible that reduced firing of GCs in the presence of inhibitory DREADDs is sufficient to produce eyeblink conditioning, albeit with a slower onset. Similarly, for VOR learning, GC-specific alterations of K-Cl cotransporters[19] or deleting CaV2.1 calcium channels[27] led to less pronounced gain changes in gain-down learning than we observed, but both of these manipulations leave significant GC signaling intact. Meanwhile, one study found that reducing GC firing by deleting the SCN2A gene was sufficient to eliminate gain down leaning for 20 min training[22], but it is not known how VOR learning is affected for more prolonged train (240 min over 4 days), as used in our study. PC-specific elimination of the AMPAR auxiliary subunit SHISA6 decreased GC-PC EPSCs by a factor of 3, but virtually eliminated VOR gain-down learning and conditioned eyeblink learning[6]. However, CF-PC synapses are also likely altered in adult SHISA6 mice, and these synapses play a crucial role in cerebellar-dependent plasticity and learning. Lastly, we show that although GC signaling is not required for VOR and VVOR, it is essential for OKR response (**Fig. 3c-d, h-i, m-n**), which was not apparent in studies in which GC signaling was suppressed but not eliminated [19, 27]. Together, these studies indicate that it is necessary to completely and selectively eliminate GC signaling to reveal the full extent of the influence of GCs on behavior.

Surprisingly, social approach and responses to social odorants were normal in GC TKO mice. What makes this even more remarkable is that GC TKO mice are ataxic, but this did not influence social preference. This was such an unexpected result that we checked for the possibility that the lack of effect might be due to remaining GC-PC signaling, which could explain the performance of mice in which CaV2.1 had been selectively eliminated from GCs[21]. However, we found that GC-PC synaptic transmission was eliminated in GC TKO mice, and intact responses to social odors and the normal social preferences could not be explained by residual synaptic transmission (**Fig. 5e-h**).

Our results also have important implications for studies of SCN2A, which is a major risk factor for ASD that is associated with heightened responses to sensory stimuli. The presence of *Scn2a* (NaV 1.2 encoding gene) in GCs, combined with the observation that heterozygous expression of NaV 1.2 in GCs leads to hypersensitive VOR[22], raised the possibility that reduced expression of NaV 1.2 in GCs could be the cause of ASD in SCN2A patients. Our results suggest that decreasing GC signaling alone is insufficient to produce social deficits associated with ASD, and it is important to directly examine social behaviors in mice with selective heterozygous loss of NaV 1.2 in GCs.

Although GCs are not required for VOR, anxiety or social approach, that does not mean that GCs in regions of the cerebellar cortex responsible for these behaviors do not have functional roles. Just as GCs are not an important regulators of baseline VOR but they are essential for gain-down VOR learning, it is likely that GCs allow the cerebellum to learn new social responses.

It is remarkable that eliminating GC signaling influences behaviors so differently than increasing GC excitability. We previously selectively eliminated the GABA_A_R δ subunit from GCs (Cb δ KO mice) to attenuate tonic GC inhibition. This, in turn, made GCs much more excitable. Whereas eliminating GC signaling had profound effects on most motor behaviors[4] and motor learning (**Fig.1** and **Fig 2**), in Cb δ KO mice motor function and motor learning were normal. We also found that anxiety was unaffected in GC TKO mice but was elevated in Cb δ KO mice. Lastly, there are no social deficits in GC TKO mice, whereas social behaviors are altered in females in Cb δ KO mice[28]. The stark behavioral differences that arise from completely silencing GC synapses, or increasing GC excitability, highlight the differences in insights that can be obtained by increasing or decreasing the influence of a particular cell type.

### Comparison of the behavioral influences of GCs and PCs

A comparison of the effects of suppressing either PC or GC outputs establishes that some cerebellum-dependent behaviors are reliant on GC signaling and some do not require GC-dependent cerebellar processing (**Table 1**). Performance on balance beam, rotarod, gait, OKR, and many spontaneous behaviors (**Fig. 4b**), as well as VOR learning and conditioned eyeblink, are all impaired similarly in GC TKO mice and PC TKO mice, indicating that these behaviors rely on intact GC signaling. Surprisingly, eliminating GC and PC signaling differentially influenced many behaviors. GC TKO mice did not exhibit a number of behavioral deficits that were observed in PC TKO mice: (1) there was no prominent overshoot in VOR (**Fig. 3h,i,j**), (2) some spontaneous behaviors were unaffected (**Fig. 4c**), (3) the distance traveled was not impaired (**Fig. 4d**) and a rapid component of movement was intact (**Extended Data Fig. 4b,c**), (4) anxiety levels did not increase (**Fig. 4e**), and (5) social approach and responses to social odorants were normal.

**Table 1.**
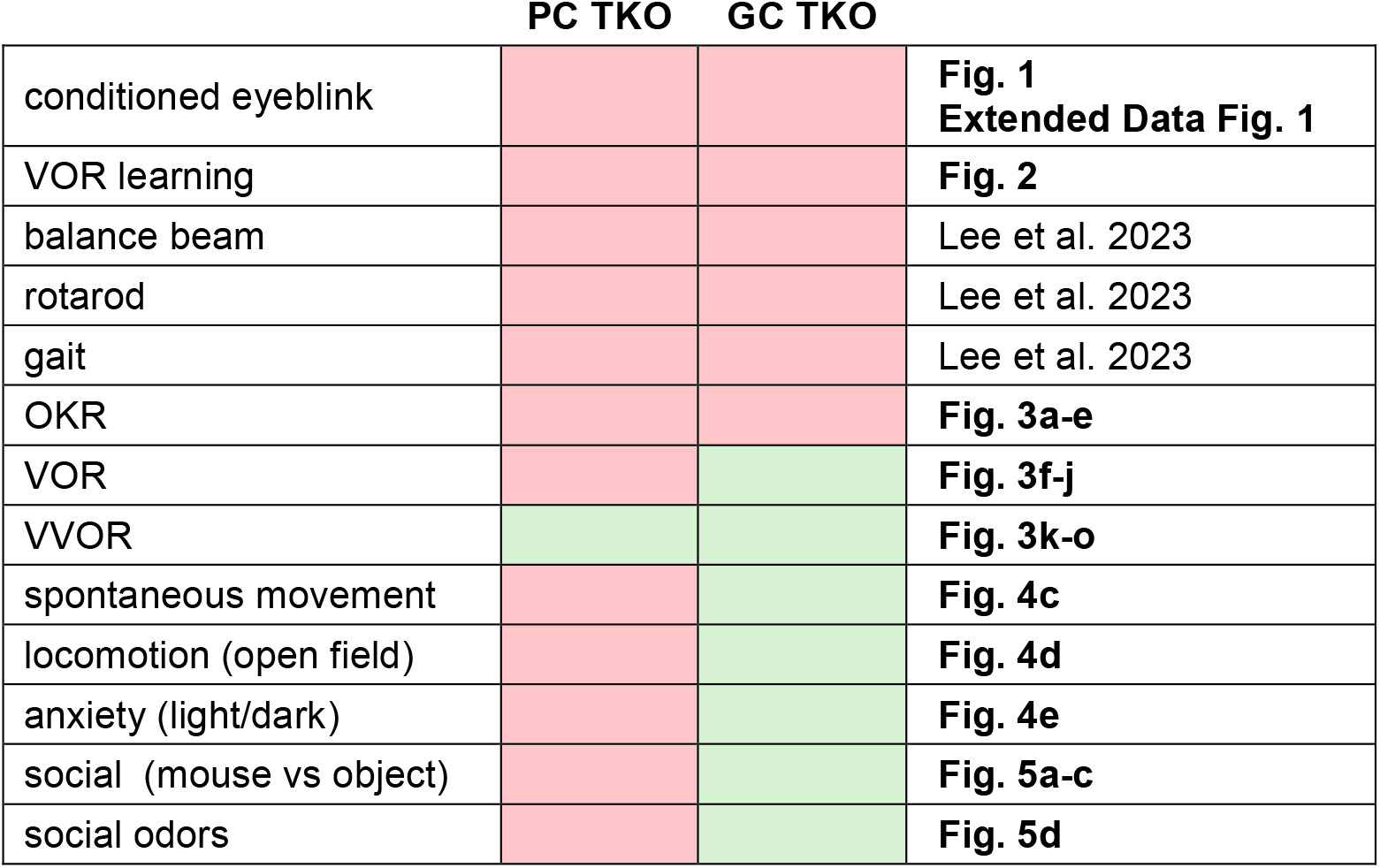
Behavioral effects of silencing the output of the cerebellar cortex (PC TKO) or the input layer (GC TKO) of the cerebellar cortex. Red indicates an impairment in behavior, green indicates that the behavior is unaffected. Some of the results have been extracted from our previous publication[4].

It is well established that particular regions of the cerebellar cortex influence specific behaviors [8, 37, 43, 44], and it was generally assumed that the same basic circuit is repeated throughout the cerebellar cortex and that different regions regulate different behaviors using the same basic circuit that adjusts the weights of GC-PC synapses to transform a pattern of mossy fiber inputs into a pattern of Purkinje cell outputs. Spontaneous PC firing is usually ignored when considering how the cerebellum influences behavior. The stark contrast between severely impaired motor learning (**Fig. 2g**) and normal social behavior (**Fig. 5c, left**) observed in GC-TKO mice clearly demonstrates that not all cerebellum-dependent behaviors are mediated by the same conventional cerebellar processing that relies on GC-PC synapses.

These findings have important implications for rescuing deficits that arise from cerebellar dysfunction. Behaviors that rely on cerebellar processing, such as controlling gait, balance, or motor learning, involve complex and dynamically regulated firing in many PCs. It is expected that it would be exceedingly difficult to correct cerebellar dysfunction that arises from alterations in PC firing dynamics. In contrast, our results suggest that VOR overshoot, elevated anxiety, and deficits in social behaviors do not involve GC-dependent processing. The most likely explanation is that the elimination of spontaneous firing in PC TKO mice leads to these deficits. This suggests that the lack of spontaneous PC firing disinhibits neurons in the cerebellar nuclei, causing increased firing in downstream regions. This raises the possibility that rescuing such behaviors is a matter of suppressing the firing of neurons in downstream regions. Interestingly, this is exactly what was previously observed in an autism mouse model, where mice exhibiting social deficits due to decreased PC firing in Crus I of the cerebellar cortex showed elevated firing in the contralateral prefrontal cortex, and suppressing firing in the prefrontal cortex rescued social behaviors [7, 8, 34]. This study suggested that ASD could be treated by elevating PC activity or by suppressing the activity of appropriate downstream regions. Our results suggest that such a therapeutic strategy was successful for social deficits, because they are not reliant on granule-cell dependent processing. Our results also suggest that a similar strategy might also be effective in treating the subset of cerebellar dependent deficits that do not arise from a deficit in GC-dependent processing, such as VOR overshoot, and elevated anxiety.

## Methods

### Mice

Animal procedures have been carried out in accordance with the NIH and Animal Care and Use committee (IACUC) guidelines, and protocols approved by the Harvard Medical Area Standing Committee on Animals. Male and female mice were used for experiments. Mice were kept on a mixed background (129Sv/SvJ and B6/C57). Mice were housed under standard conditions in groups of 2-5 animals on a 12 h light-dark cycle with food and water available *ad libitum*.

We generated GC (a6) TKO, GC (gabra6) TKO and PC (PCP2) TKO mice as we described beforele[4]. Briefly, we utilized the previously described mouse line that has floxed alleles for the conditional ablation of CaV2.1, CaV2.2, and CaV2.3. (*floxed* CaV2s line)[45]. This mouse line was crossed with *a6-cre* line (RRID: MMRRC_000196-UCD) for GC (a6) TKO, *Gabra6-cre* line (RRID:MMRRC_000213-UCD) for GC (gabra6) TKO, and *Pcp2-cre* line (Jackson laboratory, Strain# 010536) for PC (PCP2) TKO, to produce the mice with the genotype Cre(+); CaV2.1(*flox*/ *flox*), CaV2.2(*flox*/ *flox*), CaV2.3(*flox*/ *flox*). The littermate mice with the genotype Cre(−); CaV2.1(*flox*/ *flox*), CaV2.2(*flox*/ *flox*), CaV2.3(*flox*/ *flox*) were used as control mice.

### Behavioral testing

All motor behavior testing was conducted on adult male and female mice, with the experimenter blinded to the genotype. For conditioned eye blink tests, the 10-11 week old mice were used. For VOR learning and OKR/VOR/VVOR experiments, 9-10 week old mice were used. For MoSeq experiment, 11-15 week old mice were used. For open-field test, light-dark test and three-chamber test, 8-13 week old mice were used. For olfaction test, 7-34 week old mice were used. For Before behavioral testing, mice were handled by experimenter for 15 mins for two days. On the test day, mice were transferred to the behavior room and allowed to acclimate for at least 30 min. Apparatuses used for behavioral testing were cleaned with 70% ethanol between experiments.

### Conditioned eye blink test

Before the experiment, mice were implanted with a head bracket and allowed to recover for at least five days post-surgery. Then, one day of habituation is performed before the first day of eyeblink training to accustom the animals to the apparatus. During the habituation, the mouse is head-fixed for 20 min atop a motorized treadmill with six inches in diameter, rotating at 2.5 cm/sec. During the training, a white LED flash to the left eye with 550 ms duration is used as the conditioned stimulus (CS), which is co-terminated with the unconditioned stimulus (US) (periorbital air puff 2 psi for 5 ms) to the right eye. A hundred CS-US paired trials and ten CS-only trials were randomly assigned and tested with a mouse per day. The intervals between the trials are randomized between 4 to 12 seconds. The training lasts for 8-9 days. During the trials, the right eye is recorded by a high-speed IR camera (Mako U-029B; Allied Vision, Exton, PA) and a macro lens (1/2 inch, 4-12mm F/1.2, Tamron, Commack, NY.) at 280 fps. To detect eye blink events, the positions of the upper and lower lids in the recorded videos are detected by an open-source deep neural network, DeepLabCut[46]. Then, the vertical distance between the upper and lower lids is analyzed. The pneumatic and electronics necessary for the control of the air puff were based on the design of Openspritzer[47]. The CR amplitude was defined as the average eye closure value measured between 0.4 and 0.5 seconds after the LED was turned on. The CR probability is defined as the percentage of events where the CR amplitude exceeds 50%. Analyses are performed using custom MATLAB and Python code.

### OKR/VVOR/VOR and VOR training

Before the experiment, mice were implanted with a head bracket and allowed to recover for at least five days post-surgery. During the test, the mice are head-fixed on the treadmill (speed, 2.5 cm/sec) with their left eye at the center of a custom-made VOR apparatus. The apparatus is fully enclosed for light control and includes a motorized turntable powered by a servo motor (CPM-MCVC-2341S-RLS; Teknic, inc.), delivering horizontal vestibular stimuli. The cylindrical paper screen surrounding the mouse with a 25 cm distance from the left eye is used to present the visual stimuli generated by Laser Screen Beam Projector (MP-CL1A, 1920×720; Sony, inc.).

The visual stimuli are the vertical bars spaced by 7 degrees with 3-degree widths, and move back-and-forth horizontally by ± 5°. The left eye is recorded by a high-speed IR camera (Mako U-029B; Allied Vision, Exton, PA) at 200 fps with 640 × 480 pixel resolution. A multifunction I/O device (PCIe-6321; NATIONAL INSTRUMENTS CORP.) is used to monitor the table rotation and to command the servo motor and the camera. The visual stimuli, I/O device and camera were controlled by a custom-made GUI MATLAB application. An infrared LED fixed to the top of the camera is used to make corneal reflection on the left eye. The infrared LED to illuminate the left eye is positioned below the camera. One day before the experiment, the mice are habituated on the apparatus for 20 min. During the experiments, pilocarpine (4% ophthalmic drops; Patterson Vet Supply, Inc.) was applied to the left eye to limit pupil dilatation to accurately track the eye movement in darkness.

Right before the eye recording, we take pictures of the pupil of each mouse while moving the camera back-and-forth around the vertical axis of the turntable with a known angle (± 10°) to calculate the radius of pupil rotation (Rp) that is used to convert the pixel position to angular position of the eye (Stahl et al., 2000). Rp values are measured at three different pupil diameters by calculating Rp = Δ/sin(20°), where Δ is the pixel distance between the centers of the pupils, determined by their relative position to the corneal reflection in images taken at ±10°. Then, the Rp regression line is made to estimate the Rp values with various pupil diameters. The angular eye movement is calculated by the following formula: angular eye movement between t1 and t2 = arcsin [(P_t1_ − CRF _t1_)−(P _t2_ − CRF _t2_)/Rp] where P is the pupil center position, CRF is the position of corneal reflection, and Rp is the value estimated by Rp regression line for the corresponding pupil diameter.

In optokenitic reflex (OKR) test, the left eye is recorded while the different frequencies of visual stimuli (0.2, 0.5 and 1 Hz) with a fixed ± 5° amplitude are presented while the turntable is fixed. In vestibulo-ocular reflex (VOR) test, the left eye is recorded in complete darkness while the turntable is rotating back-and-forth with different frequencies (0.2, 0.5 and 1 Hz) with a fixed ± 5° amplitude. The visually guided VOR (VVOR) test is the same as the VOR test, except the fixed visual stimuli is presented. In VOR training, the visual stimuli and the turntable are rotated in phase (at the same amplitude, ± 5°) on Day 1. On Day 2-4, the visual stimuli are rotated in phase to the turntable rotation but with greater amplitudes (Day2, ± 5°; Day3, ± 7.5°; Day4, ± 10°), while the amplitude of the turntable remained ± 5°. On each training day, 6 × 10 min VOR trainings are performed, and 7 VOR tests are performed before, between, and after the trainings, to probe the effect of the trainings. In VOR training, the frequencies of the visual stimuli and the turntable are both 0.5 Hz. In VOR tests, the frequency of the turntable is 0.5 Hz.

In the acquired video, the pupil center is detected using an open-source deep neural network, DeepLabCut[46]. After the pixel velocity of the pupil center is converted to angular velocity, a low-pass filter set at twice the stimulation frequency is applied. Data exceeding 40°/sec is removed from further analysis to exclude saccadic eye movements, except for VOR response analysis, where a threshold of 60°/sec is applied. This was because PC-TKO mice exhibited non-saccadic VOR responses exceeding 40°/sec. Gain values are calculated as the ratio of the amplitude of angular eye velocity to the angular velocity of either the visual stimulus or the turntable velocity. Analyses are performed using custom MATLAB and Python code.

### Motion Sequencing

Motion Sequencing (MoSeq)-based behavioral analysis is performed as in the previous studies[28, 30, 31]. In brief, MoSeq uses unsupervised machine learning techniques to identify the number and content of behavioral syllables out of which mice compose their behavior; identifying these syllables allows each video frame of a mouse behavioral experiment to be assigned a label identifying which syllable is being expressed at any moment in time. Behavioral phenotypes that distinguish wild-type and mutant mice can be identified by comparing differences in how often individual syllables are used in each experiment. Here, individual mice are imaged for three 30-minute-long sessions using a Kinect2 depth sensor while behaving in a circular open field. These 3d imaging data are submitted to the MoSeq pipeline[30].

### Open Field Test

During this test, an animal was placed in uncovered rectangular behavior arena (30.3 cm × 45.7 cm, 30.5 cm high) and allowed to explore for 10 minutes, under dim (40 lux) lighting. The position of the mouse was tracked using custom MATLAB scripts. The travel distance and the time spent in the center zone, corresponds to the four central areas when the bottom surface is evenly divided into 16 sections, was calculated.

### Light-Dark test

The light-dark chamber (40 cm × 20 cm) consists of a light chamber (>600 lux) and a dark chamber (<10 lux). A door between two chambers allows the mice to freely explore the two chambers. At the beginning of the experiment, the mouse is placed in the dark chamber and allowed to freely navigate both chambers for 10 min with videotaping. The position of the mouse was tracked using custom MATLAB scripts. Then, the ratio of time spent in the light chamber over the dark chamber is analyzed.

### Three chamber assay

The arena consists of a clear rectangular Plexiglas box (40.5 cm wide, 60 cm long, 22 cm high) without a top cover and is divided into three equally sized compartments by two transparent walls. Each wall contains a 10.2 cm × 5.4 cm rectangular door to allow the mice to navigate between the compartments. First, mice are pre-exposed to the middle chamber for 5 minutes, with the doors to the adjacent chambers closed, to maximize the exploration time in the side chambers during the test. Then, during a 10 minute baseline session, the doors are opened, and mice are allowed to freely navigate all three chambers for 10 minutes. After that, the doors are closed again while the animal remains in the middle chamber. Then, a wire cup (10 cm in diameter) containing a social stimulus (juvenile mouse aged 15-30 days of the same sex) is placed in either the right or left chamber. Also, a wire cup containing a novel object (mouse-sized plastic toy, Schleich GmbH, Germany) is placed in the opposite chamber. The sides of social and object stimuli are randomly selected to control for preference to either side of the arena. After the stimulus placements, the doors are opened, and the animal is observed for another 10 minutes. The time spent in the social chamber and object chamber are analyzed offline using a custom-made MATLAB code.

### Olfaction test

The experiments are performed in the dark environment (<10 lux). Mice are placed in a transparent chamber (18 cm wide, 32 cm long, 13 cm high) with 2-3 mm bedding and habituated with a cotton-tipped applicator without odor for 30 min. The cotton tip is placed 2 cm above the bottom in the middle of the chamber. Then, the cotton tip with 10 µL of odor is presented three times for 2 minutes each. For each 2 min recording, the cotton tip is replaced with a new cotton tip with the same odor. During the presentation, the mice are videotaped by IR camera (EPL IR USB camera, 30 Hz frame rate). In this way, water, coconut, urine A (same sex), urine B (opposite sex) are presented. The coconut odor is 10 times diluted from the original liquid (McCormick, inc.). The urine samples are collected from unfamiliar, sex-matched adult mice aged over 7 weeks. In the recorded videos, the odor responses are manually counted by the experimenter blind to the genotype. The odor responses are defined by the sniffing behavior when mouse’s nose is facing the cotton tip and the distance to the cotton tip is less than 3 cm.

### In vivo recording

To prepare for in vivo recordings, mice (9-10 week old) are anesthetized with isoflurane (4-5% induction, 1-3% maintenance) and implanted with a custom-made titanium head bracket. A small craniotomy (0.5 – 1 mm diameter) is drilled over the Crus I (2.4 mm from the midline, −6.0 mm posterior to the bregma). The cranium above the anterior cerebellum remained exposed and is covered with Kwik-Sil, a silicone polymer, at the end of the surgery. Mice are allowed to recover from surgery for at least five days. For acclimation, mice are head-restrained on a free-moving cylindrical treadmill for at least 30 minutes the day before recording. On the day of recording, the silicone polymer covering the cranial window is removed, and a silicon probe (64 channel probes, Cambridge NeuroTech) dipped in Di-I (Vybrant Multicolour Cell Labelling Kit, Thermofisher) is inserted into the brain. In vivo recordings were performed while the mouse was in a quiescent state. Data are sampled at 20 kHz using an RHD2000 recording system (Intan Technologies) and filtered between 0.1 and 8kHz. PCs are distinguished by the presence of complex spikes. When recordings are complete, mice are perfused with PBS and 4% PFA. 100 μm coronal slices are made from the brain tissue to confirm recording location. Single units are sorted using open-source modules, Kilosort[48], SpikeInterface[49], and Phy2 (https://github.com/cortex-lab/phy). Analyses are performed using custom Python code.

### Slice preparation for electrophysiology

Male mice of 8 weeks old were used for in-vitro physiology experiments. Animals were anesthetized with ketamine / xylazine / acepromazine and transcardially perfused with warm choline-ACSF solution containing in mM: 110 Choline Cl, 2.5 KCl, 1.25 NaH_2_PO_4_, 25 NaHCO_3_, 25 glucose, 0.5 CaCl_2_, 7 MgCl_2_, 3.1 Na-Pyruvate, 11.6 Na-Ascorbate, 0.005 NBQX, and 0.0025 (R)-CPP, oxygenated with 95% O2 / 5% CO2. To prepare sagittal slices of the cerebellum, the hindbrain was first removed. A cut was made down the cerebellar midline, and the two halves of the cerebellum were glued with the medial face down to the slicing chamber. 150-200 µm thick sagittal slices were cut with a Leica 1200S vibratome in warm choline-ACSF. Slices were transferred to a standard ACSF solution containing, in mM: 127 NaCl, 2.5 KCl, 1.25 NaH_2_PO_4_, 25 NaHCO_3_, 25 glucose, 1.5 CaCl_2_, and 1 MgCl_2_ maintained at 34-35°C for 10-12 minutes and then stored at room temperature for at least 20 minutes before beginning recordings.

Whole-cell recording of PCs were obtained with borosilicate glass electrodes of 1-2 MΩ resistance. All recordings were done at 33°C with 2.5uM SR95531 (gabazine) in the bath to block inhibitory input. Visually guided recordings of PCs in Crus1 were performed while holding the cell at -65mV with an internal containing (in mM): 35 CsF, 110 CsCl, 10 Hepes, 10 EGTA and 2 QX-314, pH adjusted to 7.2 with CsOH and osmolarity adjusted to 307 mOsm/kg. Electrophysiology data was acquired using a Multiclamp 700B amplifier (Axon Instruments), digitized at 20 kHz and filtered at 4 kHz. For PF stimulation, theta glass electrodes filled with ACSF were placed in the molecular layer (∼70 um apart) and applied with brief (0.5 ms) current pulse (50 µA) for single stimulation and 20 Hz current pulses (30 µA) for train stimulation. For single stimulation experiments, evoked response in each PC was averaged from 30-40 trails in 3-4 stimulating locations.

### Statistical Analysis

All statistical tests, significance analyses, number of individual experiments (n) and other relevant information for data comparison are specified in Table S1. Statistical analysis was performed using commercial software (Graphpad Prism 9) and open-source Python library[50]. For all analyses, norm ality tests were performed first to appropriately select parametric or non-parametric methods. Significance levels are indicated as *p < 0.05, **p < 0.01, ***p < 0.005 and not significant (ns). No statistical methods were used to predetermine sample sizes. In box plots, the bottom and top of each box are the 25th and 75th percentiles of the sample, respectively. The distance between the bottom and top of each box is the interquartile range. The bottom and top error bars indicate the most extreme values within 1.5 times the interquartile range from the 25th and 75th percentiles, respectively.

## Acknowledgments

We thank Shreya Palwayi, Lillian Bergin, Hailey Goffin, and Juliana Preston for their assistance with the behavioral experiments. We thank Megan R. Carey for valuable feedback. This work was supported by grants from the NIH (R35NS097284 to W.G.R.), the William Randolph Hearst Fund to J.-H.L., and the Lefler Center at HMS to Shuting Wu. Imaging was performed in the Neurobiology Imaging Facility at Harvard Medical School.

## Author Contributions

J.-H.L. and W.G.R. conceived the experiments. J.-H.L. and C.G. established the equipment and protocols for the behavioral experiments. S.W. conducted *in vitro* electrophysiology experiment. A.N., Z.Y., J.-H.L. conducted behavioral experiments. J.-H.L. analyzed behavioral data. J.-H.L. conducted *in vivo* recording and analysis. J.-H.L. visualized all figures. J.-H.L., W.G.R. wrote the manuscript with inputs from the authors.

