## supplemental figures for "Processing reliant on granule cells is essential for motor learning but dispensable for many cerebellar-dependent behaviors"

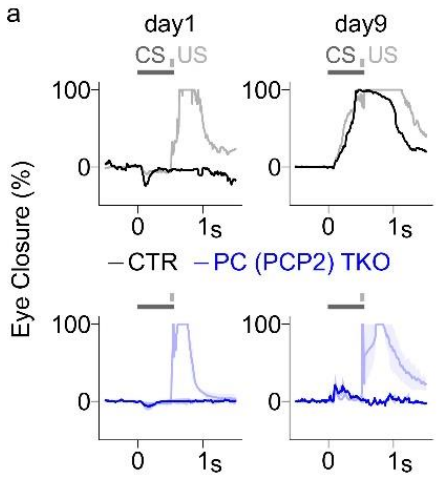


**Extended Data Fig. 1 | Responses of PC (PCP2) TKO mice in the conditioned eye blink test. a,** Average eye closures on days 1 and 9 are shown for PC (PCP2) TKO mice (n=2 mice) and littermate control mouse (n=1). CS-only trials (dark) and CS-US paired trials (faint) are plotted. The shaded area indicates SEM. Note that on day 9, we observed a response in PC (PCP2) TKO mice that resembles the short-latency response reported in a previous study [51].

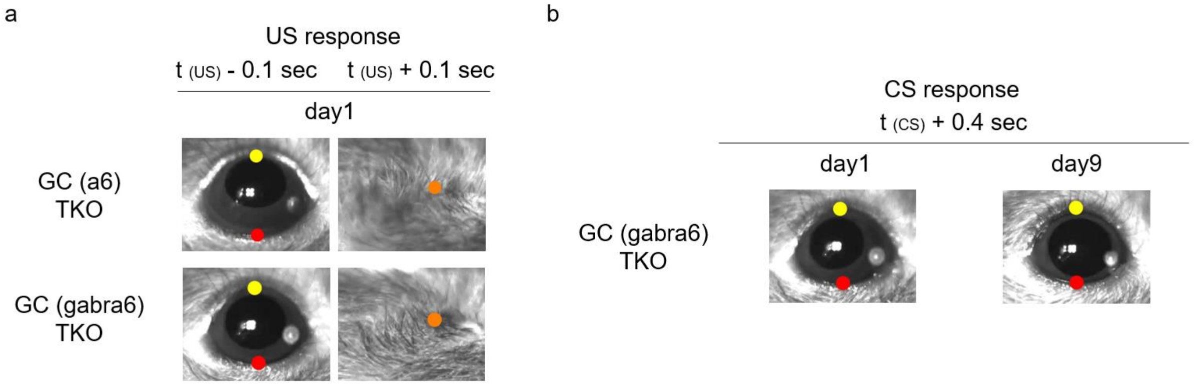


**Extended Data Fig. 2 | Examples of unconditioned responses and conditioned responses. a,** The response to the unconditioned stimulus (air-puff) is shown. The images show the eye responses at 0.1 sec before the air-puffs (*left*) and 0.1 sec after the air-puffs (*right*) on day1 for GC (a6) TKO (*top row*) and GC (gabra6) TKO mouse (*bottom row*). The time point of unconditioned stimulus is denoted as 't (US)'. **b,** Responses to the conditioned stimuli (white LED) on day 1 and day 9 of GC (gabra6) TKO mouse are shown.

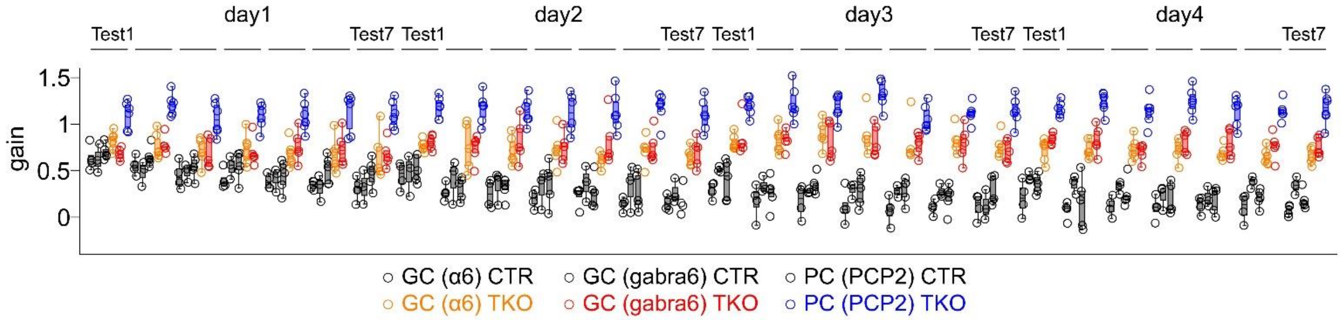


**Extended Data Fig. 3 | Collective representation of gain values during VOR learning.** The data shown in **Fig. 2g-i** are replotted for comparison, with each circle indicating a mouse.

**Supplementary Video 1 | An example of the conditioned response in the eye blink test.** The conditioned responses of the control mouse on day 1 (left) and day 8 (right) are shown. The CS (LED) was presented at 0 s, and the US (air puff) was presented at 0.5 s. The dots colored yellow, red, and orange indicate the positions of upper eyelid, lower eyelid, and closed eyelid, respectively, as automatically detected by deep-learning.

**Supplementary Video 2 | An example of pupil tracking in VOR test.** The left and right edges of the pupil (*green dot*) and the position of corneal reflection (*yellow cross*) were detected using deep-learning. The calculations of gain values were processed by custom MATLAB code.

**Supplementary Video 3 | Video of behavioral syllables.** The video recorded by 3D depth camera shows the syllables shown in **Fig. 4b** and **4c**. These crowd movies were generated by Moseq2 analysis pipeline[30], by overlaying 20 individual syllable movies from different mice exhibiting the corresponding syllables. The red dots indicate the moments when syllables occur.

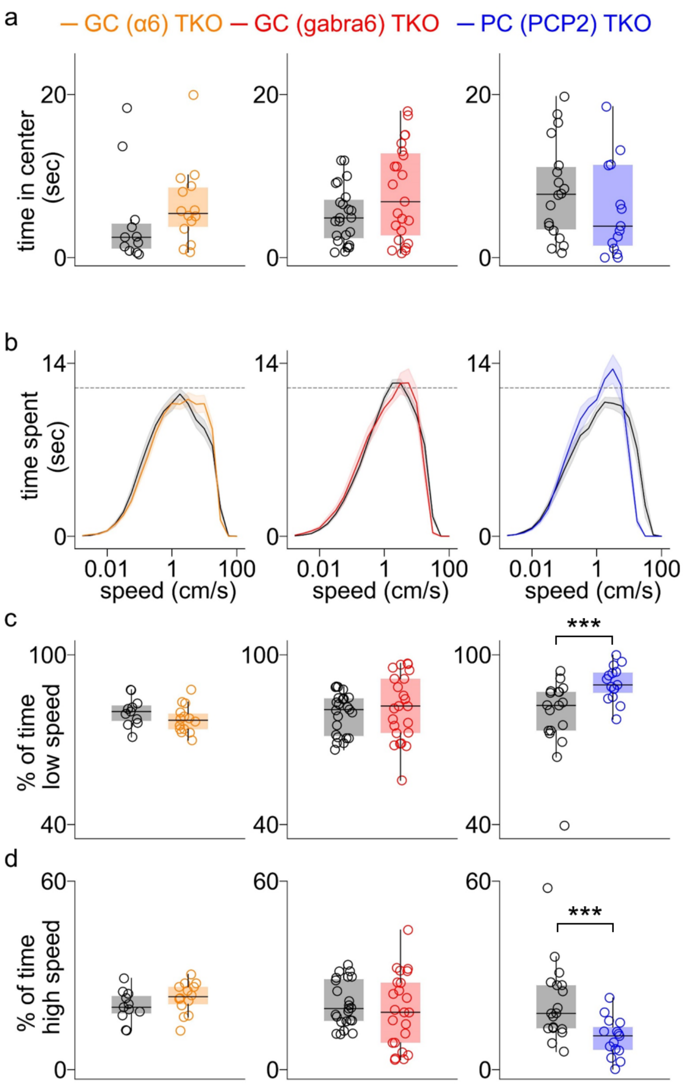


**Extended Data Fig. 4 | Properties of GC (α6) TKO, GC (gabra6) TKO and PC (PCP2) TKO mice in open field test.** **a,** The time spent in the center zone of TKO (*colored*) and control (*black*) mice are shown. Each circle indicates a mouse. **b-d,** A more detailed analysis of the speed of mice in the open field tests of **Fig. 4d** is provided. **b.** The time spent moving at different speeds is shown on a log scale. Plots are averages for all mice and shaded regions are ±SEM. The distribution of speeds is very similar for all genotypes with the exception of PC (PCP2) TKO mice that speed more time moving slowly and less time moving rapidly. **c.** The percentages of time spent at low speed (0-5 cm/s) is compared for each types of TKO and its own controls. **d.** As in c, but for high speed (20-100 cm/s). PC (PCP2) TKO mice spend significantly more time moving at low speed and less time moving at high speed.

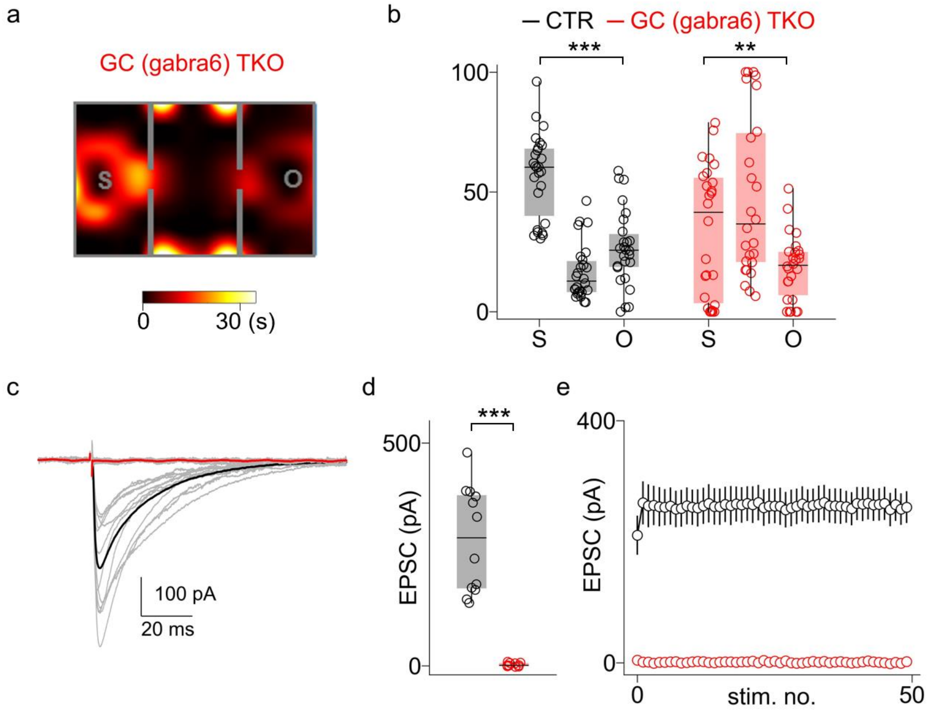


**Extended Data Fig. 5 | Social behavior of GC (gabra6) TKO mice. a,** As in **Fig. 5a** but for GC (gabra6) TKO mice. **b,** As in **Fig. 5c**, the time spent in social chamber (S), middle chamber and object chamber (O) during sociability test are shown for GC (a6) TKO (*orange*) and control (*black*) mice. Each circle indicates a mouse. GC (gabra6) TKO mice spend more time in the chamber with the mouse than with the object, as for GC (α6) TKO mice. However, they also spend more time in the corners of the center chamber. This is likely a result of the properties of the GABRA6-Cre line (see text). **c,d,e,** Same as in **Fig 5e, 5f, 5h** but for GC (gabra6) TKO mice.

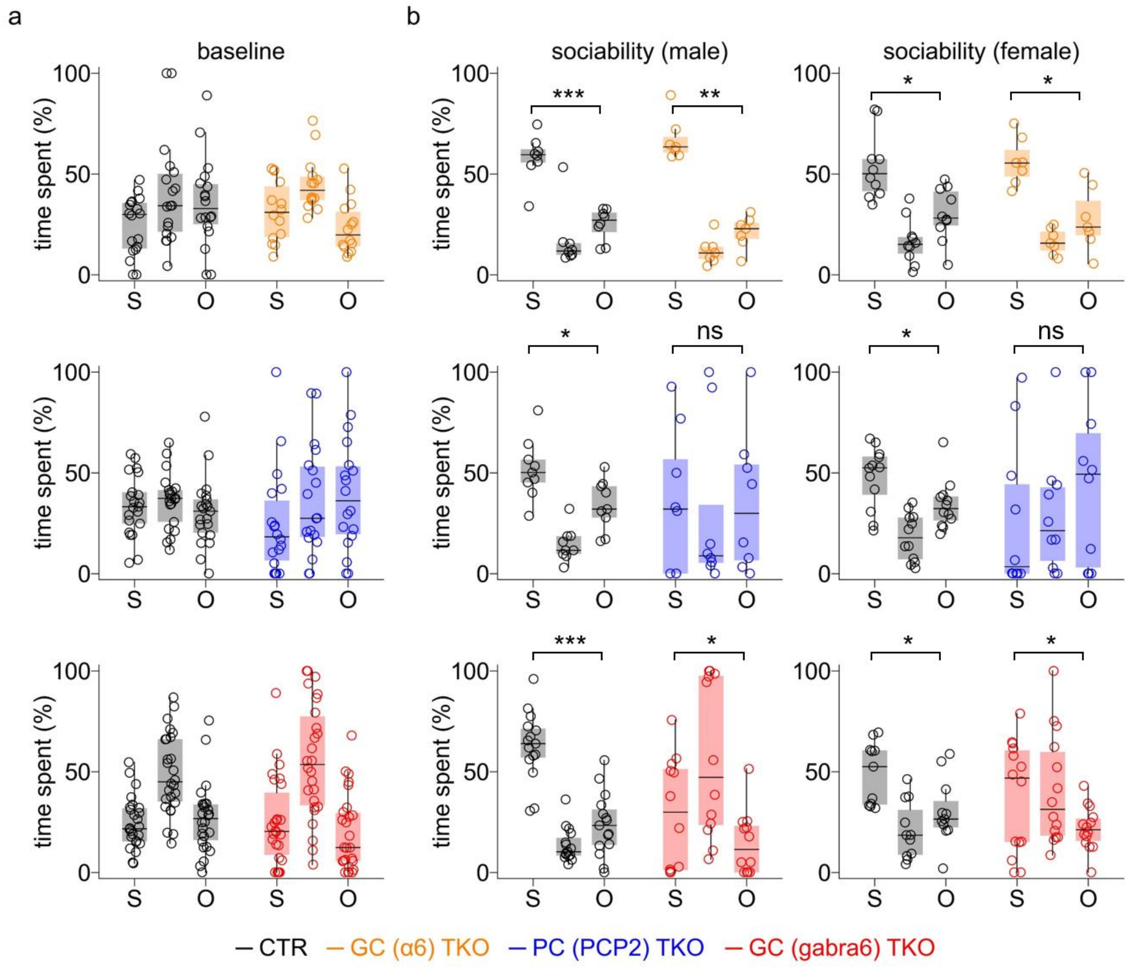


**Extended Data Fig. 6 | Properties of social behaviors of GC (a6) TKO, GC (gabra6) TKO and PC (PCP2) TKO mice. a,** The times spent in social chamber (S), middle chamber and object chamber (O) during the 10 min baseline are shown for control (*black*), GC (a6) TKO (*orange*), PC (PCP2) TKO (*blue*) and GC (gabra6) TKO (*red*) mice. Each circle indicates a mouse. **b,** Same as in **Fig. 5c** and **Extended Data Fig. 5b**, but males (*left*) and females (*right*) are plotted separately.
